# Agonistic anti-CD40 converts Tregs into Type 1 effectors within the tumor micro-environment

**DOI:** 10.1101/2022.10.17.512537

**Authors:** Vivien Maltez, Charu Arora, Rina Sor, Qiaoshi Lian, Robert H. Vonderheide, Ronald N. Germain, Katelyn T. Byrne

**Author notes:** Correspondence (R.N.G.), (K.T.B.). These authors contributed equally (Ronald N. Germain, Katelyn T. Byrne).

## Abstract

Multiple cell types, molecules, and processes contribute to inhibition of anti-tumor effector responses, often frustrating effective immunotherapy. Among these, Foxp3+ CD4+ cells (Tregs) are well-recognized to play an immunosuppressive role in the tumor microenvironment. The first clinically successful checkpoint inhibitor, anti-CTLA-4 antibody, may deplete Tregs at least in part by antibody-dependent cellular cytotoxicity (ADCC), but this effect is unreliable in mice, including in a genetically engineered mouse model of pancreatic ductal adenocarcinoma (PDAC). In contrast, agonistic CD40 antibody, which serves as an effective therapy, is associated with notable Treg disappearance in the PDAC model. The mechanism of CD40-mediated Treg loss is poorly understood, as Tregs are CD40-negative. Here we have explored the mechanistic basis for the loss of Foxp3 T cells upon anti-CD40 treatment and find, using tissue-level multiplex immunostaining and orthogonal dissociated cell analyses, that Tregs are not depleted but converted into interferon-*γ* (IFN-*γ*) producing, Type I CD4+ T effector cells. This process depends on IL-12 and IFN-*γ* signaling evoked by action of the anti-CD40 antibody on dendritic cells (DCs), especially BATF3-dependent cDC1s. These findings provide insight into a previously unappreciated mechanism of CD40 agonism as a potent anti-tumor intervention that promotes the re-programming of Tregs into tumor-reactive CD4+ effector T cells, markedly augmenting the anti-tumor response.

---

Immunotherapy has revolutionized the field of medical oncology, yet most patients are resistant to current interventions. The immunobiology of the tumor microenvironment (TME) is intrinsically suppressive, often resulting in subpar immunological responses to immune checkpoint blockade (ICB) that are insufficient to drive or maintain clinical responses in patients^1^. Pancreatic ductal adenocarcinoma (PDAC) is a canonical example of immunotherapy-refractory disease, with >98% of patients resistant to ICB.

Recently, we and others have contributed to a growing understanding of the critical roles that immunotherapy-induced CD4+ T cells play in mediating tumor rejection^2–5^. CD4+ T cells may directly kill tumor cells via novel cytotoxic mechanisms^6^, or play a ‘helper’ or instructor role, mediating anti-tumor immunity indirectly via tumoricidal macrophages^7^, CD8+ T cells^4^, humoral immunity^8^, and/or stromal cell crosstalk^9^. Using the *Kras*^*G12D*^*Trp53*^*R172H*^*Pdx-1-Cre* YFP (KPCY) mouse models of PDAC^10,11^, our work revealed the potency of CD4+ T cells in rejecting established tumors after agonistic anti-CD40 (αCD40) therapy combined with ICB^12^. Targeting CD40 via agonistic antibodies mimics natural CD40 ligand engagement and enables dendritic cell (DC) licensing and maturation sufficient to drive T cell-mediated tumor rejection^13–15^. Recent reports reveal that DCs are required for optimal generation of *both* CD8+ and CD4+ T cell responses against tumors^12,16^, highlighting αCD40 as an intervention that ‘pushes the gas pedal’ on anti-tumor immunity. Concomitant with the generation of effector CD4+ T cell responses after CD40 stimulation, we have also reported a loss of immunosuppressive Foxp3+ regulatory CD4+ T cells (Tregs) in the tumor site beginning shortly after αCD40, alone or in combination with other therapeutic interventions including chemotherapy and/or ICB^12,17,18^. The precipitous drop in the Treg compartment removes a second barrier to effector T cells in the TME by ‘releasing the brakes.’

The discovery that anti-CD40 results in loss of Foxp3+ CD4+ T cells in the TME was surprising because the target of the anti-CD40 antibody is not expressed by Tregs in contrast to the checkpoint inhibitor target CTLA-4, making Treg removal after αCD40 by processes such as ADCC unlikely. Here we have studied the mechanism involved in the loss of Foxp3+ Tregs upon αCD40 treatment. For a model system, we employed an immunologically ‘hot’ PDAC-derived clonal cell line to induce flank tumors in C57Bl/6 mice^11^. As a baseline control, we repeated prior therapy studies, giving mice with established tumors αCD40 (clone FGK4.5, ‘F’) and dual ICB (αPD-1 and αCTLA-4, ‘PC’)^12^. As with the mixed clone parental tumor^12^, treatment with P every 3 days, C on day 0, 3, and 6, and a single dose of F on day 3 led to tumor regression and increased survival in mice treated with FPC, effects not seen with F or PC alone (Extended Data Figure 1A-B).

With these functional data in hand, we sought to use tissue multiplex imaging to better understand the changes induced in the TME by these therapeutic interventions, especially the loss of Tregs. We chose this imaging approach to complement the more widely used flow cytometric analysis of extracted cells because of concerns about quantitative extraction of activated T cells from tumors^19^ and also because this method would provide insight into the spatial organization of cell populations within the TME. At 48 hours after anti-CD40 administration in FPC treated mice, imaging of tumor tissue sections showed a clear reduction in Tregs (Figure 1A-B), with the remaining Tregs residing at the tumor periphery (Figure 1C). In contrast, Foxp3-CD4+ as well as CD8+ T cells retained a more homogenous distribution across the tumor site (Figure 1C), revealing that the Treg compartment was uniquely affected in terms of number and location upon combination antibody treatment.

**Figure 1.**
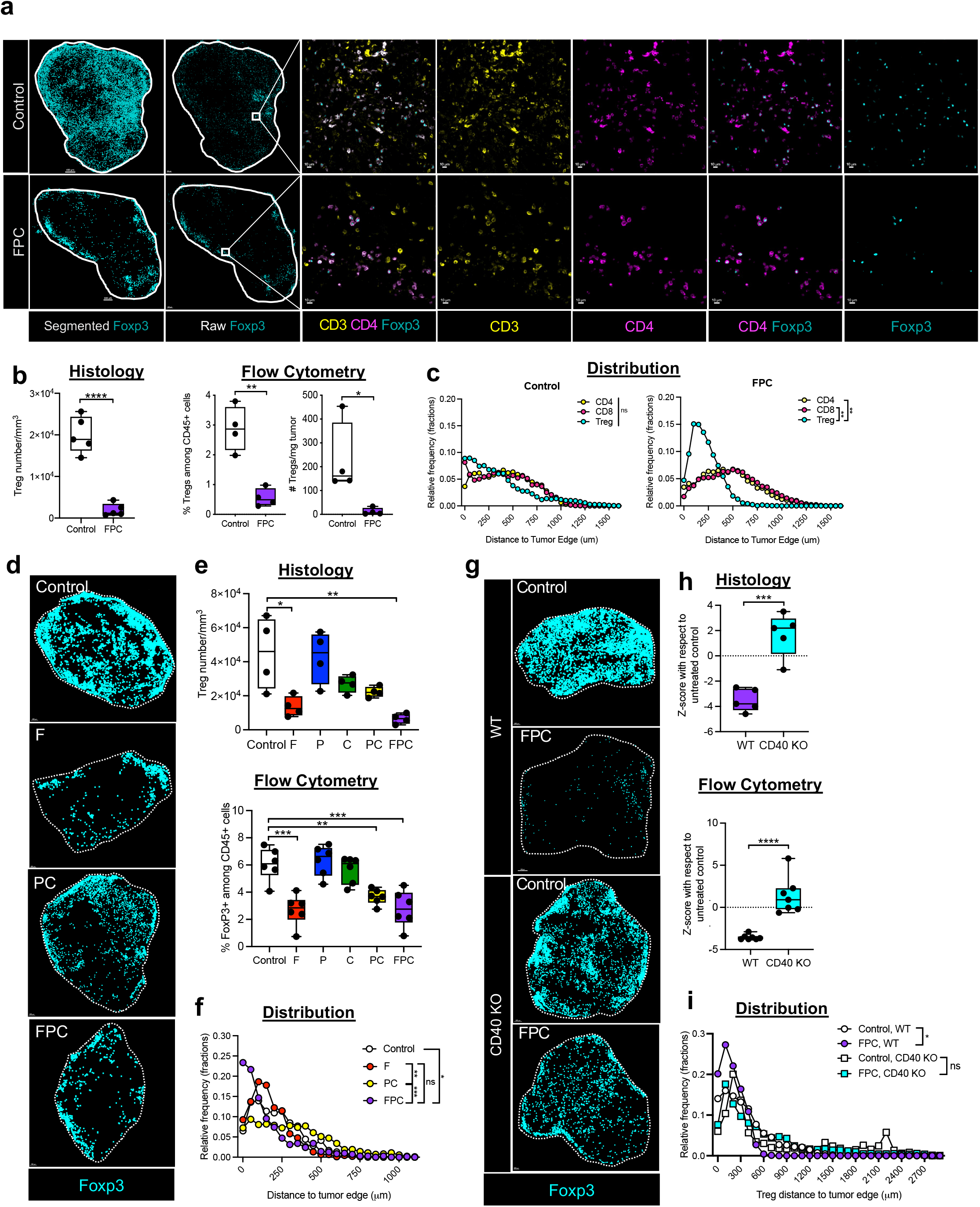
Agonistic CD40 therapy reduces intratumoral Tregs independently of anti-CTLA-4 treatment. Wild-type (WT) C57BL/6 mice were injected with KPCY T cell high tumor clone 2838c3 and treated with anti-PD-1 and anti-CTLA-4 (PC) on days 0 and a single dose of agonistic anti-CD40 (F) on day 3, alone or in combination (FPC). Tumors were harvested on day 5 for analysis. (A-B) Tumors were stained for Foxp3 expression (A) with a representative example of segmented dots (which will be used for all subsequent figures) and raw marker expression. The zoom inset shows the raw expression of the indicated markers. Total Tregs were quantified in (B) by tissue immunostaining (left) or flow cytometric analysis among live, CD45+ cells or total Tregs per mg of tumor (right). Total CD4+ or CD8+ T cell or Treg distribution was quantified in (C) by relative frequency distributions with bins set every 100 μm. (D-F) Mice were treated as in 1A with individual treatment components as indicated (D) and quantified by tissue immunostaining (left) or flow cytometry (right) (E) as in 1B, or for intratumor distribution (F). (G-I) Wild-type or CD40 knock out (KO) mice were treated as in 1A (G), and the normalized fold-change in Tregs within the TME after therapy was calculated for both tissue immunostaining (left) and flow cytometry (right) (H) as in 1C or distribution as in 1D (I). Data are representative of 2-5 independent experiments with n=5-10 mice per group; each symbol represents an individual mouse in box and whisker plots, horizontal lines indicate the median, bars show interquartile range, and whiskers indicate the range. Scale bars for A are: Control, 400 μm; FPC, 300 μm, zoom insets, 15μm; for D: Control, 200 μm; PC and FPC, 300 μm; F alone, 500 μm; for G: CD40 KO Control, 500 μm, all others are 200 μm. Dotted line indicates tumor edge (A, D, G). Analysis by unpaired T-test of z-score normalized to control group (B, H), or one-way ANOVA with Tukey’s post-test and mean difference calculations to account for effect size (C, E, F, I). * indicates p<0.05, ** indicates p<0.01, *** indicates p<0.001, **** indicates p<0.0001. For distribution plots (C, F, I): * indicates mean difference of >50 um, ** >100 um, ***>150 um, ****>200 um

Intratumoral Treg reduction has been reported in some tumors as a result of αCTLA-4-mediated ADCC^20,21^, so we next assessed the contribution of each component of FPC to Treg loss and distribution in the TME. Alterations in the intratumoral Treg compartment occurred after treatment with αCD40 alone or in combination with ICB (Figure 1D-E). Furthermore, Tregs were predominantly located at the tumor edge only after treatments including αCD40 (Figure 1F). In contrast, αCTLA-4 alone or in combination with αPD-1 did not consistently reduce the absolute Treg numbers or influence the Treg distribution in the tumor site as compared to control treated mice (Figure 1D-F)^12^. To further explore the requirement of agonistic αCD40 in mediating Treg reduction, we repeated our analysis after FPC therapy in wild-type (WT) and global CD40 knockout (KO) mice. In the absence of functional CD40, the reduction in Tregs was no longer seen after treatment with FPC, whether assessed by imaging or flow cytometry (Figure 1G-H). Further, in these KO mice FPC treatment no longer led to a concentration of residual Tregs at the tumor edge (Figure 1I).

Given that T cells do not express CD40^22^ we hypothesized that a professional antigen presenting cell (APC), such as a DC – a major target of αCD40 antibody therapy^23^ – may play a role in mediating intratumoral Treg changes after FPC. To determine if major histocompatibility complex (MHC) II-expressing cells that act as APCs to CD4+ T cells participate in the Treg reduction observed after therapy, WT mice were treated with an MHC II blocking antibody beginning two days before the start of FPC. MHC II blockade abrogated both the Treg loss and tumor edge localization induced by FPC therapy (Extended Data Figure 2A-C), implicating both antigen presentation by CD40-expressing APCs and antigen receptor engagement by T cells in αCD40-mediated Treg loss. To further explore the role of DCs in Treg reduction, we took advantage of mice lacking expression of Batf3, as conventional Type 1 (cDC1) subsets fail to develop in the absence of this transcription factor^24^. cDC1s are the primary APC for CD8+ T cells but are also a major APC for CD4+ T cells and the locus for delivery of CD4+ T cell ‘help’ to CD8+ T cells^16,25,26^. Tumor-bearing WT or Batf3 KO mice were treated with FPC and assessed for intratumoral Tregs after therapy. In the absence of cDC1s, Tregs were not reduced by treatments containing anti-CD40 (Figure 2A-C). To examine if these results might be related to generation of a T cell low (FPC-resistant) PDAC TME because of the absence of cross-presenting DCs at the time of tumorigenesis^11^, these studies were repeated in tumor-bearing CD11c diphtheria toxin receptor (DTR) mice. In these animals, multiple subpopulations of DCs can be specifically depleted via DT administration two days prior to FPC treatment, after the TME is established. DC ablation using DT also abrogated FPC-induced Treg loss and concentration at the tumor periphery (Extended Data Figure 2D-F), indicating the strict requirement for DCs in CD40-mediated Treg loss.

**Figure 2.**
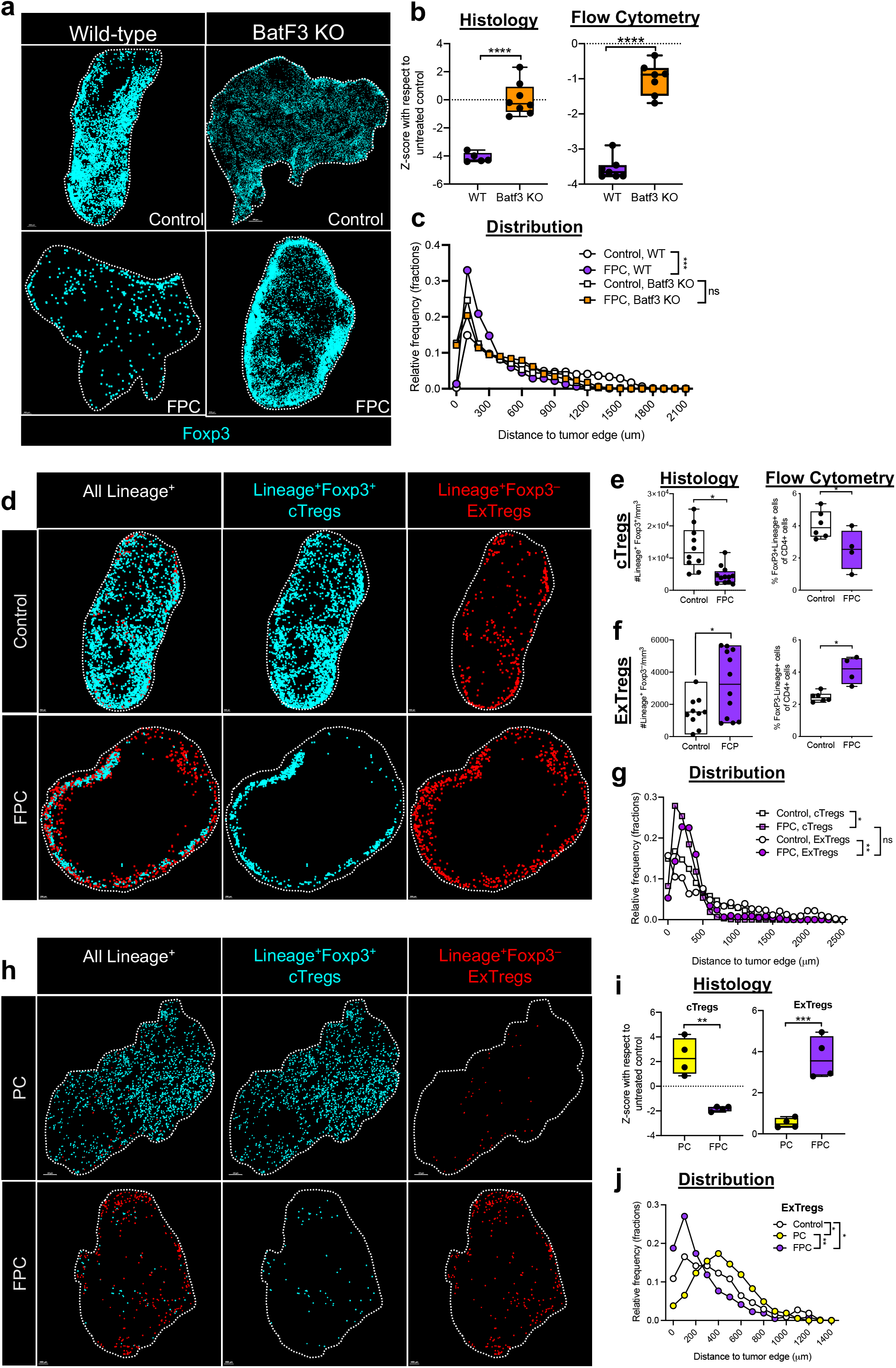
Anti-CD40 antibody induces Treg conversion via dendritic cells. (A-C) Tumor-bearing WT or Batf3 knockout (KO) mice were treated and tumors were analyzed by tissue immunostaining (A) and analyzed as in Fig 1A-D for Treg proportions (B) and distribution (C). (D-J) Experiments were performed in Foxp3 Lineage mice as described in Fig 1 with the addition of tamoxifen day 8-10 to label Tregs with the lineage marker, tdTomato. (D-G) Tumors were analyzed by tissue immunostaining (D) for expression of Lineage trace marker and Foxp3, with Lineage+Foxp3+ conventional cTregs quantified in (E) and Lineage+Foxp3-ExTregs quantified in (F) by both histological and flow cytometric analysis. (G) Distribution analysis of indicated Treg subsets was performed as in 1C. (H-J) Comparison of tumors from Foxp3 Lineage mice treated with PC +/-F by tissue immunostaining (H) for proportions of cTregs or ExTregs (I) and distribution of ExTregs (J). Data representative of 2-5 independent experiments with n=3-12 mice per group; each symbol represents a single mouse, horizontal lines indicate the median, bars show interquartile range, and whiskers indicate the range. Scale bars for A are: WT Control, 200 μm; Batf3KO Control, 500 μm; both FPC treated, 400 μm; for D and H: 200 μm for all. Dotted line indicates tumor edge (A, D, H). Analysis by unpaired T-test of z-score normalized to control group (B, E, F, I), or one-way ANOVA with Tukey’s post-test and mean difference calculations to account for effect size (C, G, J). For p values, * indicates p<0.05, ** indicates p<0.01, *** indicates p<0.001, **** indicates p<0.0001. For distribution plot (C, G, J): * indicates mean difference of >50 um, ** >100 um, ***>150 um, *ns* indicates not significant.

What might antigen-presenting cells be doing in response to αCD40 that causes the loss of Foxp3+ CD4+ T cells? One possibility was some mechanism of induced cell death, but another possibility was loss of Foxp3 expression without cell death. In the latter case, we would see a numerical loss of Tregs as assessed by dual Foxp3 and CD4 staining without the T cells actually dying. To address this issue, we first assessed the expression of the apoptotic marker, cleaved caspase 3 (CC3), in the Treg compartment 48 hours after treatment with αCD40 and found that the proportion of CC3+ Tregs was equivalent with or without FPC treatment (Extended Data Figure 2G).

To more directly determine if Tregs were losing expression of Foxp3 in the PDAC TME after αCD40, we utilized *Foxp3*^*eGFP-Cre-ERT2*^ x *Gt(ROSA)26Sor*^*tm(CAG-tdTomato)/Hze*^ (R26tdTomato) mice to conduct lineage tracing experiments^27^. In these mice (hereafter referred to as Foxp3 Lineage mice), Foxp3-expressing cells are permanently labeled by tdTomato expression upon treatment with tamoxifen, while real-time expression of Foxp3 is reflected by GFP expression. Tumors were implanted and allowed to establish in Foxp3 Lineage mice, at which point mice were treated with tamoxifen for three days to irreversibly label all Foxp3-expressing Tregs. Mice were then rested for two days before treatment with FPC followed by harvesting and assessment of the tumors 48 hours after αCD40 administration as before. Conventional Tregs (cTregs) were identified as Tregs that expressed Foxp3 and tdTomato (Lineage+Foxp3+), while ExTregs were identified as cells that retained tdTomato expression but lacked Foxp3 expression (Lineage+Foxp3-) (Figure 2D-F). Upon assessing the PDAC TME in Foxp3 Lineage mice after treatment with FPC, the cTreg compartment was observed to be significantly reduced in FPC-treated vs. control mice (Figure 2D-E), as expected. However, the ExTreg compartment showed a significant increase in the TME of FPC-treated mice, both in absolute number and proportion, as compared to the ExTreg compartment in the TME of control mice (Figure 2D, F and Extended Data Figure 2H). The proportion of CC3+ cTregs (and ExTregs) was similar, regardless of treatment with FPC (Extended Data Figure 2I), indicating that the change in frequency and number within the Treg compartments of control and FPC-treated mice was not a result of increased cell death specifically within the cTreg subset. Both cTregs and ExTregs were concentrated at the edge of the tumor in FPC treated mice (Figure 2G).

Because αCD40 was uniquely required for loss of Foxp3+ CD4+ T cells in this model, we assessed the development of ExTregs in PC vs. FPC treated mice. Treatment with PC was insufficient to induce a robust ExTreg population in the tumor site (Figure 2H-I) or their polarized localization at the tumor edge (Figure 2J), revealing a requirement for αCD40 in mediating cTreg → ExTreg conversion that paralleled the inverse loss of Foxp3+ Tregs.

DCs play a unique role in bridging innate and adaptive immune responses via both direct and indirect mechanisms of communication with T cells, and the IL-12/IFN-γ cytokine axis plays a major role in this cellular crosstalk, particularly with regards to CD4+ T cells^28^. Prior work indicated that strong inflammation accompanied by substantial IL-12 and IFN-*γ* production in the gut could lead to partial Foxp3 loss by Tregs and acquisition of a more Type 1 effector-like phenotype^29^. This led us to examine the role of these cytokines in the formation of ExTregs in the TME after CD40 agonism. Loss of host expression of either IL-12p40 or IFN-γ prevented FPC-induced control of tumor growth and animal survival, consistent with a large literature showing that the Type 1 inflammatory axis contributes to immune control of cancer^30,31^ (Extended Data Figure 3A-B). Tumors from both IL-12p40 KO and IFN-γ KO mice also retained the Foxp3 Treg (cTreg) compartment after FPC therapy (Figure 3A-B). To bypass the impact of an altered baseline TME due to systemic absence of either of these two cytokines, WT mice were treated with blocking antibodies targeting IL-12p40 or IFN-γ beginning one day before the start of FPC therapy. Again, the αCD40-mediated reduction and polarized location of residual Foxp3+ Tregs after FPC treatment was largely abrogated when either IL-12p40 or IFN-γ was neutralized (Figure 3C-E) revealing the requirement for signaling via the IL-12/IFN-γ cytokine axis to mediate tumor rejection and concomitant cTreg reduction in the PDAC TME after therapy. Furthermore, IL-12 or IFN-γ blockade in combination with FPC showed that perturbation of this cytokine axis blunted the conversion into ExTregs and their concentration at the tumor edge (Figure 3F-H). Thus, Foxp3+ Tregs convert to an ExTreg population in the PDAC TME after αCD40 stimulation in an IL-12/IFN-γ dependent manner.

**Figure 3.**
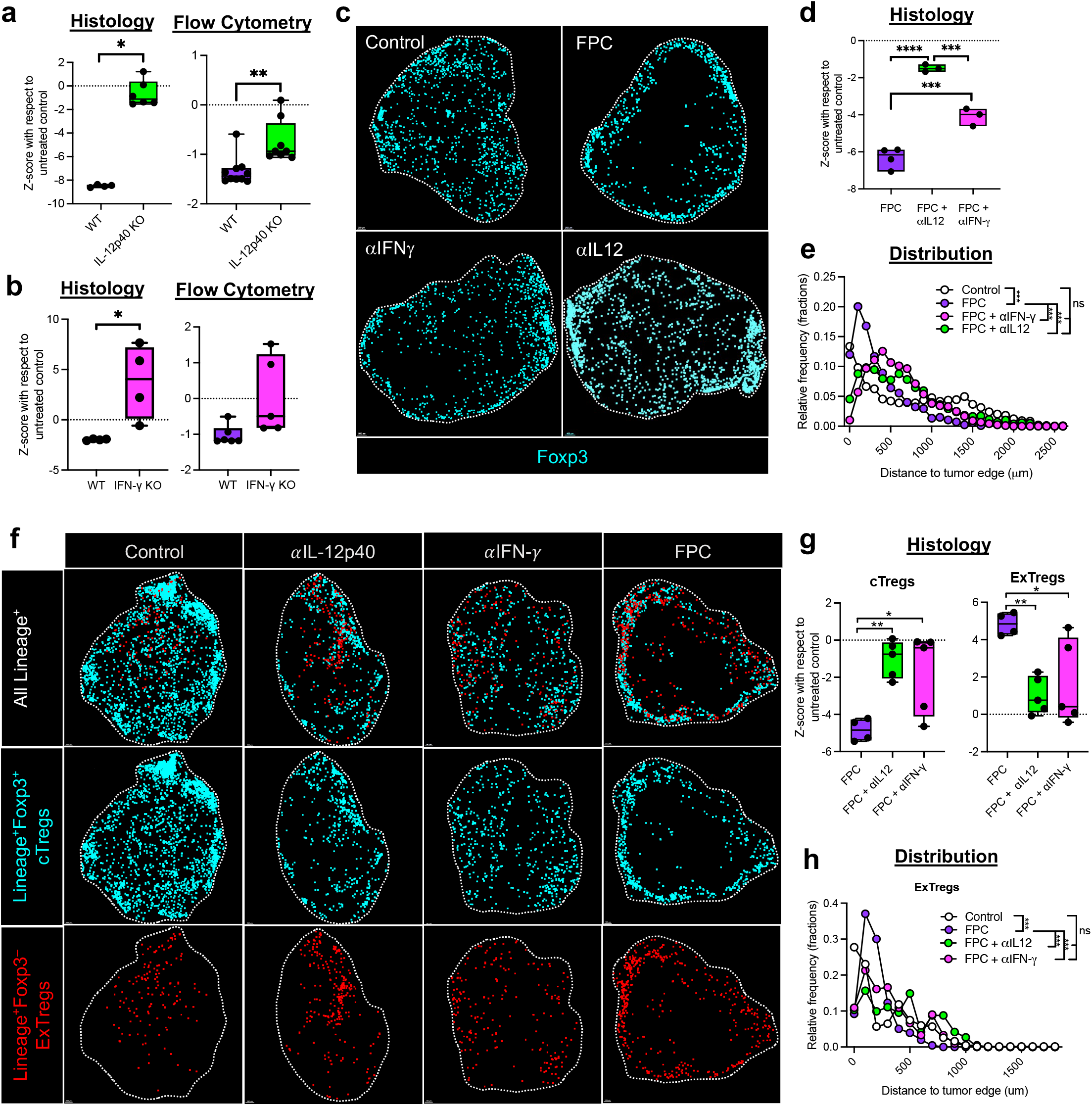
The IL-12/IFN-*γ* axis mediates Treg conversion after anti-CD40 therapy. Mice were treated as described in Fig. 1A. IL-12p40 (A) or IFN-γ (B) knockout (KO) tumor-bearing mice were treated and tumor were analyzed as in Fig 1A for Treg proportions by tissue immunostaining (left) or flow cytometric (right) analyses. (C-E) WT mice bearing established tumors were treated with IL-12p40 or IFN-γ blockade two days before the start of therapy, and tumors were analyzed by tissue immunostaining (C) for Treg proportions (D) and distribution (E). (F-G) Foxp3 Lineage mice were treated +/-FPC, or FPC + IL-12p40 or IFN-γ blocking antibodies, started 2 days before FPC treatment, analyzed by tissue immunostaining (F) for cTregs or ExTregs (G, left and right, respectively) and distribution of ExTregs (H). Data representative of 2-5 independent experiments with n=3-5 mice per group; each symbol represents an individual mouse, horizontal lines indicate the median, bars show interquartile range, and whiskers indicate the range. Scale bars for C are: Control and FPC, 200 μm; anti-IL12p40, 400 μm; anti-IFNg, 300 μm; for F are: Control and anti-IL12p40, 200 μm; anti-IFN-γ, 150 μm; FPC, 100 μm. Dotted line indicates tumor edge (C, F). Analysis by unpaired T-test of z-score normalized to control group (A, B, D), one-way ANOVA with Tukey’s post-test and mean difference calculations to account for effect size (E, H), or one-way ANOVA with Tukey’s post-test (G). For p values, * indicates p<0.05, ** indicates p<0.01, *** indicates p<0.001, **** indicates p<0.0001, *ns* indicates not significant. For distribution plot (E): *** indicates mean difference >150 um, *ns* indicates not significant.

The preceding studies revealed that anti-CD40 reduces the population of Foxp3-expressing CD4+ T cells by altering Foxp3 levels in these cells, not by eliminating them from the TME, as a proportion of CD25+ cTregs was retained even after αCD40 therapy (Extended Data Figure 4A). As Foxp3 has been shown to be critical to the suppressive activity of CD4+ Tregs, this αCD40-induced loss of Foxp3 expression can limit negative effects of Tregs in the TME, but is this all that the treatment does? The intestinal studies cited above suggest that Tregs exposed to an inflammatory environment can gain Type 1 effector function^29^, which could make a substantial contribution to effective tumor immunotherapy. We therefore interrogated the phenotype and function of both the cTreg and ExTreg subpopulations in the PDAC TME after treatment with FPC. Using a panel of antibodies to transcription factors, we observed that the proportion of Tbet+Foxp3+ cells, but not TCF1+ or Blimp1+ Tregs, was slightly increased after FPC treatment (Extended Data Figure 4B). This was confirmed using a Tbet/Foxp3 reporter mouse, in which we observed an increased proportion of Tbet+Foxp3+ double positive cells after αCD40 treatment (Extended Data Figure 4C-D). In FPC treated mice, cTreg and ExTregs were also both enriched for Tbet expression (Extended Data Figure 4E). The gain in Tbet expression by both Tregs that retained measurable Foxp3 and those that lost it (but were lineage marked) is consistent with prior studies showing gain of Tbet expression in cells concomitant with partial loss of Foxp3 expression in Type 1 inflammatory conditions^29^, indicative of a potential Th1-like Treg population as well as conversion to ExTregs.

Although Foxp3 is a master regulator of the Treg compartment, Helios is well established as a marker of Treg cell stability and suppressive functionality^32–34^. To interrogate the stability of the Treg compartments, Helios expression was assessed in tumors from Foxp3 Lineage tracing mice. A substantial proportion of cTregs expressed Helios in both control and FPC-treated conditions (Figure 4A-B), suggesting they are conventional thymus-derived suppressive Tregs^34^. In contrast, the ExTreg population lost Helios expression under both control and FPC-treated conditions (Figure 4A-B), suggesting loss of suppressive functionality. Additionally, the reduction in Helios expression in FPC-treated cTregs compared to the control cTregs is congruent with our data indicating an overall shift towards a Th1-like phenotype even within the cTreg compartment in FPC-treated tumors (Figure 4B).

**Figure 4.**
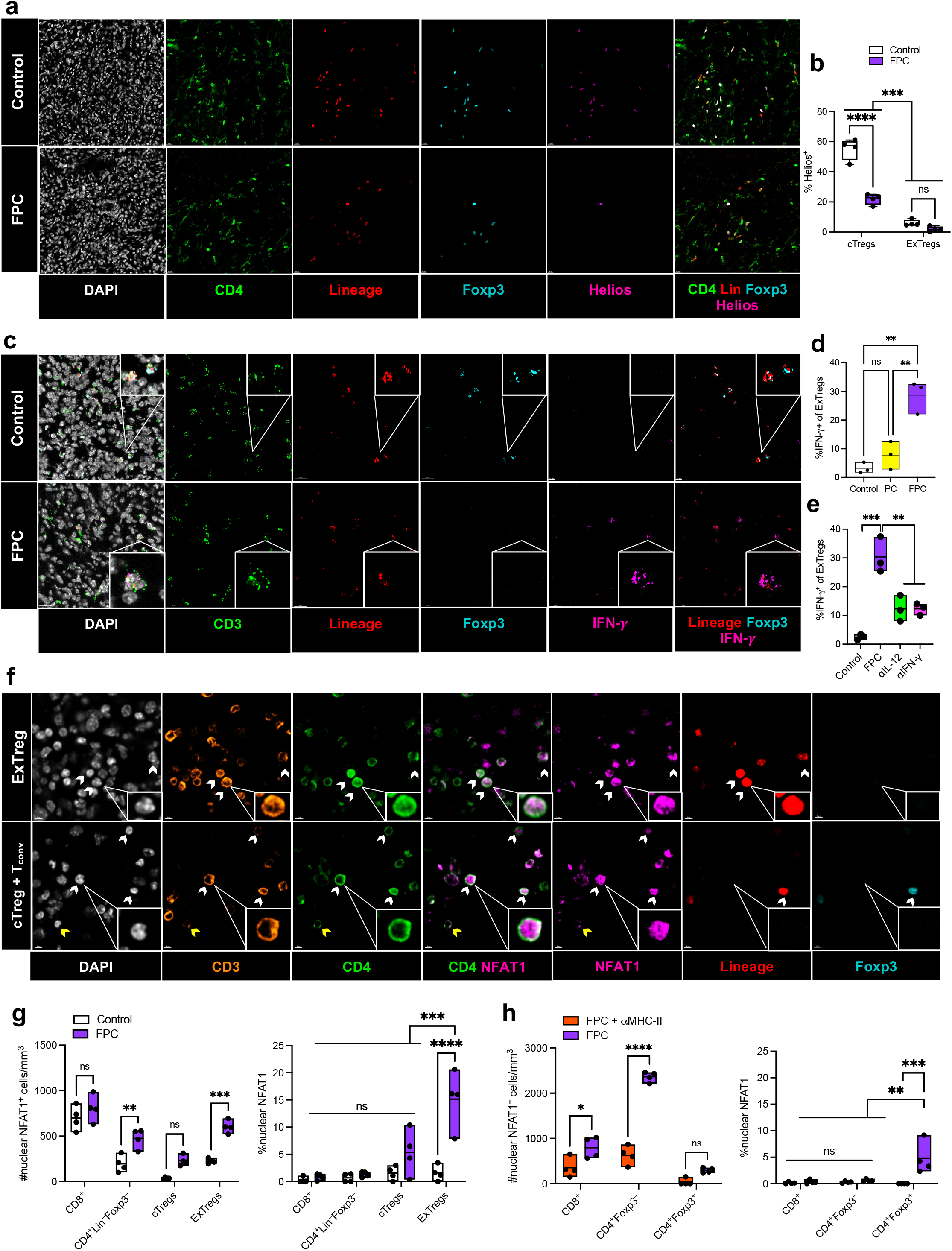
Tregs acquire a Th1 IFN-*γ* producing phenotype after αCD40 therapy. (A-G) Foxp3 Lineage mice were treated +/-FPC (A-G) or treated FPC +/-MHC-II blockade as in Ext Data Figure 2a (H) and tumors assessed *in situ* for various markers. (A) Representative immunostaining for Helios in control and FPC treated tumors, with (B) quantifications. (C) Representative RNAscope imaging with (D-E) quantifications of cTregs and ExTregs among the indicated treatments. RNA probes for indicated markers are shown. (F) Representative tumor immunostaining for nuclear NFAT1 (white arrows) and membrane NFAT1 (yellow arrows) expression within a variety of T cell populations, with examples for ExTregs (F, upper), cTregs and effector CD4 (non-Treg) T cells (T_conv_) (F, lower). (G-H) Quantification of nuclear NFAT1 by number (left) and percentages (right). Data representative of 2-3 independent experiments with n=3-5 mice per group; each symbol represents an individual mouse, bars indicate range and horizontal line indicates the mean (B, D-E, G-H). Scale bars for A are: 15 μm; for C are: 20 μm; for F: 5 μm. Analysis by unpaired T-test (B), one-way ANOVA with Tukey’s post-test (D-E), or two-way ANOVA with Tukey’s post-test (G-H). For p values, ** indicates p<0.01, *** indicates p<0.001, **** indicates p>0.0001, *ns* indicates not significant.

Tbet expression by bona fide suppressive cTregs with Foxp3 co-expression has been reported by multiple laboratories^35–37^. The functional activity of ExTregs that express Tbet and have lost Foxp3 is unclear, as the elimination of Foxp3 expression vs. competition for transcriptional control could result in a very different phenotype^38^. To examine this question, we performed RNAScope on control or FPC-treated tumors to probe for IFN-γ transcripts in ExTregs directly in the TME (Figure 4C), as IFN-γ is a canonical Type 1 effector cytokine. Both the proportion and absolute numbers of ExTregs producing IFN-γ transcripts in situ without exogenous stimulation were significantly increased in the TME of FPC-treated mice as compared to control treated mice (Figure 4C-D, Extended Data Figure 5A-B). Furthermore, PC-treated mice displayed no significant increase in IFN-γ+ cells within the small population of ExTregs present in the PDAC TME (Figure 4D and Extended Data Figure 5A). Consistent with the observation that blockade of the IL-12/IFN-γ axis disrupted αCD40-induced changes in the Treg compartment, the proportion of IFN-γ producing ExTregs was also reduced in the tumors of mice treated with IL-12 or IFN-γ blocking antibodies (Figure 4E and Extended Data Figure 5B). There have been multiple reports of exhausted T cells making IFN-γ mRNA despite defects in IFN-*γ* protein production^39–42^. To complement our RNAScope findings, we therefore examined ExTregs for IFN-γ protein by flow cytometry (Extended Data Figure 5C). Despite the limitations of dissociating these cells from the TME, our results were consistent with the RNAscope findings of an IFN-γ producing ExTreg population.

Given the requirement for intact IFN-γ signaling in the generation of ExTregs, we hypothesized that ExTregs may have altered transcriptional profiles as a result of feed-forward IFN-γ stimulation. Indeed, phosphorylated STAT1 (pSTAT1) (indicative of IFN signaling^43,44^) showed increased expression in the ExTreg subset in the TME from FPC-treated mice (Extended Data Figure 5D-E). Accordingly, both IL-12 and IFN-γ blockade abrogated pSTAT1 signaling in the ExTreg compartment (Extended Data Figure 5F-G). Given our previous finding that the Type I IFN receptor (IFNAR) is not required for FPC efficacy^5^, these data also place IFN-γ as the major driver of pSTAT1 signaling after FPC treatment.

The observation that the pSTAT1/IFN-γ signaling signature was increased in both cTregs and ExTregs after FPC therapy suggests that Tregs were in proximity to, and able to sense, cognate antigen. It is historically difficult to interrogate polyclonal T cell responses in poorly immunogenic tumors without the use of a model antigen. However, sustained cytosolic Ca^2+^ is a well characterized surrogate for T cell activation, resulting in the nuclear translocation of nuclear factor of activated T cells (NFAT1)^45^. This translocation occurs rapidly within three minutes of antigen recognition but is reversed within the next 40 minutes^46^. Two-photon intravital imaging studies have previously used this rapid rheostat nature of NFAT1 translocation as a proxy for antigen sensing and T cell activation^46–48^. We first assessed our ability to visualize nuclear NFAT1 in an OT-II adoptive transfer system, where we could clearly identify nuclear NFAT1 signal within the OT-II compartment in an antigen-specific manner within the TME (Extended Data Figure 5H-I). To our knowledge, this is the first use of this technique in fixed frozen tissues. Applying this to the PDAC TME, we distinctly visualized nuclear NFAT1 within multiple T cell compartments (Figure 4F and Extended Data Figure 5J). Strikingly, although there was observable nuclear NFAT1 within all T cell compartments (Figure 4G, left), the proportion of ExTregs with nuclear NFAT1 expression (∼15%) was significantly higher than all other T cell subsets, regardless of treatment status (Figure 4G, right). Consistent with our expectation that nuclear NFAT1 reflected TCR signaling upon cognate antigen engagement, MHC-II blockade blunted nuclear translocation of NFAT1 in T cells (Figure 4H). Combined, these results indicate that nuclear NFAT1 localization can be used within fixed tissues *in situ*, is a reliable indicator of T cell activation in polyclonal T cell systems, and further supports the premise that ExTregs convert to fully functional effectors that are antigen-sensing and capable of producing the key anti-tumor cytokine IFN-γ. The high proportion of nuclear NFAT1+ ExTregs suggests that ExTregs are detecting more relevant antigen and/or have the highest avidity for such antigen. This indicates that ExTregs, when removed from the suppressive Treg pool, represent a very substantial reduction in the suppressive capacity of the Treg subset within the TME as well as a source of high levels of IFN-γ.

As with many immune cell types, Tregs exist on a spectrum, not a fixed endpoint. Treg plasticity is highly dependent on the local cytokine milieu^49,50^. In the tumor site, the suppressive Treg state is established and maintained by unstable DC-Treg interactions^51– 53^, begging the question: can Tregs be reprogrammed quickly and efficiently in the TME? Here, we show that treatment of tumor-bearing animals with agonistic αCD40 antibody promoted intratumoral Treg conversion to a Tbet+ IFN-γ producing population of “ExTreg” CD4+ T cells within 48 hours, with a concomitant increase in Tbet+ cTregs. These changes in the Treg compartment were dependent on host expression of both CD40, IL-12, and IFN-γ, and required DCs, but was independent of anti-CTLA-4, and resulted in a spatial reorganization of the tumor-immune microenvironment.

The conversion of classical Tregs to effector CD4 T cells producing IFN-γ – known for its critical role in immune-mediated tumor regression – was coupled with loss of residual Tregs from the core of the tumor (Figure 1). However, non-Treg CD4 and CD8 T cells were retained in this central region, resulting in the ratio of effector:suppressor T cells in the tumor center skewing towards effectors after αCD40 therapy and likely a major contributing factor in tumor rejection. Even among the cTregs that remained after treatment there was a higher proportion with Th1 qualities, including increased Tbet and CD25 expression with a concomitant loss in Helios expression (Figure 4 and Extended Data Figure 4). Treg plasticity towards a Th1 phenotype has previously been reported^50^ and in some cases, reprogrammed Tregs retain some Foxp3 expression but gain additional proinflammatory capacities^35–37^. In our studies, the majority of Tregs in the TME underwent complete conversion to an ExTreg phenotype after agonistic αCD40, losing Foxp3 and Helios expression but acquiring Tbet and IFN-γ expression.

The rapidity of ExTreg generation after agonistic αCD40 administration suggests that the Foxp3 Treg cell lineage could be a previously underappreciated reprogrammable tool to improve immunotherapies. Rather than assuming an accumulation of Tregs is irreversibly linked to a poor prognostic outcome, Treg plasticity might instead be leveraged to improve patient outcomes. The power to change the allegiance of a pro-tumoral, immunosuppressive cell subset, even if fleeting, could provide critical early modifications in the TME that dictate therapeutic responsiveness.

## Acknowledgements

We would like to thank the members of the Byrne lab, Vonderheide lab, and Germain lab for helpful conversations related to this work. We are especially grateful to Andrea Radtke for advice and for establishing and refining some of the methods used in this paper. This research was supported by the Intramural Research Program of NIAID, NIH (R.N.G.), NIH grants R01CA229803 (R.H.V.), a Bench-to-Bedside supplement (R.H.V. and R.N.G.), NIGMS PRAT Fellowship Fi2GM133442 (V.I.M.), and the Parker Institute for Cancer Immunotherapy (R.H.V. and K.T.B.).

## Methods

### Mice

Mice were bred and maintained under specific pathogen-free conditions at an American Association for the Accreditation of Laboratory Animal Care (AAALAC)-accredited animal facilities within the NIAID or the University of Pennsylvania. Mice were housed in accordance with the procedures outlined in the NIH Guide for the Care and Use of Laboratory Animals (NIAID Protocol # LISB-4E), or in compliance with the procedures that were reviewed and approved by the Institutional Animal Care and Use Committee of the University of Pennsylvania (Protocol #804666). Unless otherwise stated, sex and age-matched littermates (6-12 weeks of age, both sexes) were used for individual experiments. The following strains were purchased from Jackson Laboratories (Bar Harbor, ME): C57BL/6 (cat# 00664), Batf3 KO (B6.129S(C)-*Batf3*^*tm1Kmm*^/J) (cat# 013755)^24^, CD40 KO (cat# 002928)^54^, IFN-*γ* KO (cat# 002287)^55^, IFNgR KO (cat# 003288)^56^, IL-12p40 KO (cat# 002693)^57^, CD11c-DTR (cat# 004509)^58^, Foxp3-DTR^59^, Foxp3CreERT2 (Foxp3^tm9(EGFP/cre/ERT2)Ayr^/J) (cat# 016961)^60^, C57BL/6J-*Rag1*^*em10Lutzy*^*/J* (cat# 034159), RCL-tdTomato (B6.Cg-*Gt(ROSA)26Sor*^*tm9(CAG-tdTomato)Hze*^/J (cat# 007909)^61^ and GREAT (cat# 017581)^62^ mice. C57BL/6J.Ly5a mice were purchased from the NIAID-Taconic exchange platform: OT-II:Rag1 [KO] (Taconic #4234), Tbet-ZsGreen[Tg] Tg(Rorc-E2-Crimson)Tg(Foxp3-RFP) (Taconic #008509) and referenced as Tbet/Foxp3 reporter mice in the text. Foxp3CreERT2 (Foxp3^tm9(EGFP/cre/ERT2)Ayr^/J) and RCL-tdTomato (B6.Cg-*Gt(ROSA)26Sor*^*tm9(CAG-tdTomato)Hze*^/J were crossed at both the NIAID and University of Pennsylvania facilities and only male F1 offspring (‘Foxp3 Lineage mice’) were used in experiments.

### *In vivo* reagents

Tamoxifen (Sigma-Aldrich, Cat#: 10540-29-1) was dissolved in corn oil at a concentration of 20 mg/ml and male Foxp3CreERT2^+^–R26tdTomato^+/-^ animals were administered 100 µl of tamoxifen emulsion (75 mg tamoxifen/kg) or corn oil control daily on days 6-8 post tumor implantation^60^. Animals were then treated with isotype or monoclonal antibody therapies as described. Diphtheria toxin (DT) was administered at 8ng/g every 48hours for CD11c-DT receptor mice^58^ or two doses at 50ng/kg 48 hours apart for Foxp3 DT receptor mice^59^.

Mice were treated with αPD-1/αCTLA-4/αCD40 as previously described^12^. Briefly, αPD-1 (RMP1-14; BioXcell; 200 μg/dose) was injected intraperitoneally (i.p.) on days 0, 3, 6, 9, 12, 15 and αCTLA-4 (9H10; BioXcell; 200 μg/dose) on days 0, 3, and 6, with a single dose of agonistic αCD40 (FGK4.5; BioXcell; 100 μg) on day 3^12^. For isotype controls, rat IgG2a (2A3; BioXcell; 200 μg) was used. All antibodies were endotoxin free.

For MHC-II blockade, mice were i.p. injected with a single 1 mg/kg dose of αMHC-II (M5/114; BioXcell, Cat # BE0108) 12 hours prior to agonistic αCD40 administration. Mice were harvested 24 hours after αCD40 (day 4 post therapy start). For isotype controls, rat IgG2a (2A3; BioXcell; 200 μg) was used. All antibodies were endotoxin free.

For IL-12p40 and IFN-*γ* blockades, mice were i.p. injected with 200 μg/dose of αIL12p40 (C17.8, BioXcell, Cat # BE0051) or αIFN-*γ* (XMG1.2, BioXcell, Cat # BE0055) one day prior to starting therapy, 12 hours before agonistic αCD40, and 24 hours after agonistic αCD40 (days -1, 2.5, and 4). For isotype controls, rat IgG2a (2A3; BioXcell; 200 μg) was used. Mice were harvested 48 hours after agonistic αCD40 (day 5 post therapy start). All antibodies were endotoxin free.

### Implantation of tumor cell clones

Kras^LSL-G12D/+^; Trp53^LSL-R172H/+^; Pdx1-Cre; Rosa26^YFP/YFP^ 2838c3 tumor cells were generated and used as previously described^11^. Cells were cultured in DMEM (high glucose without sodium pyruvate) with 10% FBS (Gibco) and glutamine (2mM) and harvested when confluent. Cells were dissociated into single cells with 0.25% trypsin (Gibco), washed with serum-free Dulbecco’s Modified Eagle’s medium (DMEM) twice, and counted in preparation for subcutaneous implantation. 2.5×10^5^ tumor cells were implanted subcutaneously, with viability >92% for each experiment. This cell line was examined by the Infectious Microbe PCR Amplification Test (IMPACT) and authenticated to be free of contamination by the Research Animal Diagnostic Laboratory (RADIL) at the University of Missouri.

### OT-II and MC38

Mice were subcutaneously injected with 5 ×10^5^ MC38 (WT) in the left flank and 5 ×10^5^ MC38-OVA (ovalbumin) in the right flank. Nine days post tumor implantation differentiated OT-II cells were transferred intravenously. In brief, naïve CD4 T cells were isolated from OVA-specific CD4 T cell receptor transgenic (OT-II) mouse spleens using the MACS kit from Miltenyi (cat# 130-104-453). Cells were then differentiated ex vivo (R&D, CellXVivo, Cat #: CDK018) for 3 days and 1.5 million cells were transferred into tumor-bearing Rag1 KO recipient mice. Tumors were harvested 3 days post OT-II transfer and imaged for NFAT1 nuclear translocation.

### Subcutaneous tumor growth, regression, and animal survival assessment

Tumor sizes were measured every 2-3 days for tumor growth assessment experiments. Tumor length and width were measured with calipers and tumor volumes were then calculated as length*width^2^/2. Tumor volumes of 500 mm^3^ were used as an endpoint for survival analysis. Tumor regressions and waterfall plots were calculated using the initial tumor size at the start of treatment to tumor size 21 days later.

### Tissue section preparation, processing, and immunostaining

At indicated time points, mice were euthanized and tumors were quickly harvested and fixed for 14-16 hr at 4&C in BD Cytofix/Cytoperm (BD Bioscience, Cat #: 554722) diluted 1:4 in PBS. Tumors were washed 3x in PBS (5 min per wash), carefully trimmed of fat using a stereo dissection microscope and fine forceps and dehydrated for 24 hr in a 30% sucrose solution made in 0.1 M PBS. Tumors were then embedded in optimal cutting temperature (O.C.T.) compound (Sakura Finetek, Cat #: 50-363-579), frozen on dry ice, and stored at –80&C. 18-50 µm tumor sections were prepared using a cryostat (Leica) equipped with a Surgipath DB80LX blade (Leica, Cat #: 14035843497). Cryochamber and specimen cooling was set to –17&C.

Tissue sections were adhered to Superfrost Plus microscopy slides (VWR, Cat #: 48311-703), blocked and permeabilized using 0.3% Triton X-100 with Fc block for 1 hr at room temperature (22&C), and washed in PBS. Tissue sections were next incubated with directly conjugated antibodies diluted in PBS for either 15 hr at 4&C or using the PELCO BioWave Pro-36500-230 microwave in conjunction with a PELCO SteadyTemp Pro-50062 thermoelectric recirculating chiller (Ted Pella). Briefly, a 2-1-2-1-2-1-2-1-2 program was used for immunolabeling, where “2” denotes 2 min at 100 W and “1” denotes 1 min at 0 W. This program was run twice for primary antibody labeling and once for secondary antibody labeling as previously described^63^. After washing 3x in PBS (5 min per wash) at 22&C, samples were mounted in Fluoromount-G (SouthernBiotech, Cat #: 0100-01), which was allowed to cure for a minimum of 14 hr at 22&C. All imaging was performed using No. 1.5 coverglass (VWR, Cat#: 48393-241). Combinations of the following organic fluorophores were used for immunostaining: Brilliant Violet 421, Alexa Fluor 488, Alexa Fluor 532, Alexa Fluor 555, eFluor 570, Alexa Fluor 594, AAT Bioquest iF594, Alexa Fluor 647, eFluor 660, and Alexa Fluor 700.

### RNAScope

Tissues were prepared and stained following the ACD biotechne user manual for RNAscope HiPlex Assay (Document # 324100-UM, Chapters 3-4, Fresh Frozen). Briefly, tumors were harvested and immediately placed into OCT and frozen fresh on dry ice. Tissue sections were cut as described above. Slides were stored until use at -80 &C, then immediately fixed in 4% PFA for 1 hour at RT. Slides were dehydrated in sequential ethanol steps (50%, 70%, 100%), and were then ready for staining using the ACD EZ-Batch Slide Holder (321716), Humidity Control Tray (310012), and HybEZ II Oven (321710) system. HiPlex probes (supplied at 50X concentration) for CD3e (314721-T1, Alexa Fluor 488), tdTomato (317041-T2, ATTO 550), Foxp3 (432611-T3, ATTO 647N), and IFNg (311391-T4, Alexa Fluor 750) were hybridized for 2 hours in the oven at 40 &C. Slides were then washed, and the probes sequentially amplified using the HiPlex8 Detection Kit (Cat #324110). Samples were counterstained with DAPI and mounted in Fluoromount-G (SouthernBiotech, Cat #: 0100-01). All imaging was performed using No. 1.5 coverglass (VWR, Cat#: 48393-241).

### Laser scanning confocal microscopy

Immunostained images were acquired using an upright Leica TCS SP8 X spectral detection system (Leica) equipped with a pulsed white light laser, 4 Gallium-Arsenide Phosphide (GaAsP) Hybrid Detectors (HyDs), 1 photomultiplier tube (PMT), 40x (NA = 1.3) and 20x (NA = 0.75) oil immersion objective lenses, and a motorized stage. For tissue sections (18-50 µm), images were acquired using the 20x objection with 1.5 zoom, z step size of 0.5-1.5 µm, and detector bit-depth of 12. RNAscope images were acquired using an inverted Leica Stellaris, which is optimized for far-red laser excitation and emission detection above 700 nm. In all experiments, image acquisition was controlled using LAS X software.

### Image processing, segmentation, and analysis

Image files generated in LAS X software were converted into ‘‘.ims’’ files in Imaris software (Bitplane) and subjected to a 1 pixel Gaussian filter to reduce noise. Image segmentation was performed in Imaris using the ‘‘Surface Object Creation’’ module, which employs a seeded region growing, k-means, and watershed algorithm to define individual cells. Segmentation artifacts were excluded using a combination of sphericity and volume thresholds, as well as manual correction. Following cell segmentation and surface creation, the mean or summed voxel fluorescence intensity values per channel were assessed in Imaris. In certain instances, these fluorescence distributions were used to selectively visualize T cells with specific phenotypes by creating discrete thresholds using the ‘‘filter’’ tool. For spatial statistics, the “shortest distance to object” function was used in Imaris which can automatically calculate the distance between two objects (such as shortest distance from a defined Treg to the tumor edge).

### Flow cytometry of murine tumor samples

At 24 or 48 hours after αCD40/ICB treatment, mice were sacrificed and tumors prepared for single cell suspension as previously described^17^. Briefly, tumors were washed with PBS, then minced and incubated for 45 minutes in 1mg/mL collagenase XI with protease inhibitor at 37°C and filtered through a 70 μM cell strainer with cold PBS supplemented with 0.5% BSA and 2mM ethylenediaminetetraacetic acid (EDTA). Cells were counted with Beckman Coulter Counter Z2 and stained with Live/Dead Fixable Aqua Dead Cell Stain Kit (Thermo Fisher Scientific). Cell surface molecules were assessed by incubating single cell suspensions from tissues with primary fluorophore-conjugated antibodies on ice for 45 minutes in PBS with 0.5% bovine serum albumin and 2mM EDTA. For intracellular cytokine quantification, single cell suspensions were incubated for 4hours at 37°C with PMA/ionomycin (Sigma) and incubated with GolgiPlug (Brefeldin A) and GolgiStop (Monensin) (BD Biosciences). Intracellular staining was performed using the Fixation/Permeabilization kit (Thermo Fisher Scientific) according to the manufacturer’s instructions. Antibodies used in flow analysis are described in the Supplementary information. Flow cytometric analysis was performed on a LSR II or Fortessa flow cytometer (BD Biosciences) and analyzed using FlowJo software (Treestar).

### Statistical Analysis

Statistical analysis of multiple comparisons was performed using one-way ANOVA with Tukey’s HSD post-test, and comparisons between just two groups were performed using Students’ unpaired t test. Significance of overall survival was determined via Kaplan-Meier analysis with log-rank analysis and tumor growth kinetics were analyzed with two-way ANOVA with mixed effects modeling. All statistical analyses were performed with Graphpad Prism 6 and 7 (GraphPad). Error bars show standard deviation (SD) or standard error of the mean (SEM) shown as indicated in legend, and p<0.05 was considered statistically significant. * indicates p<0.05, ** p<0.01, *** p< 0.001, and **** p< 0.0001 unless otherwise indicated. *ns* denotes not significant. Significance of cell distribution (distance to edge) was first acquired using the Imaris “shortest distance to edge” calculation. These data were then analyzed using a one-way ANOVA with Tukey’s HSD post-test and mean difference analysis to account for the large size of the data sets, and for the graphs, * indicates mean difference of >50 um, ** >100 um, ***>150 um, ****>200 um in addition to having a p value of <0.001.

---

**Extended Data Figure 1.**
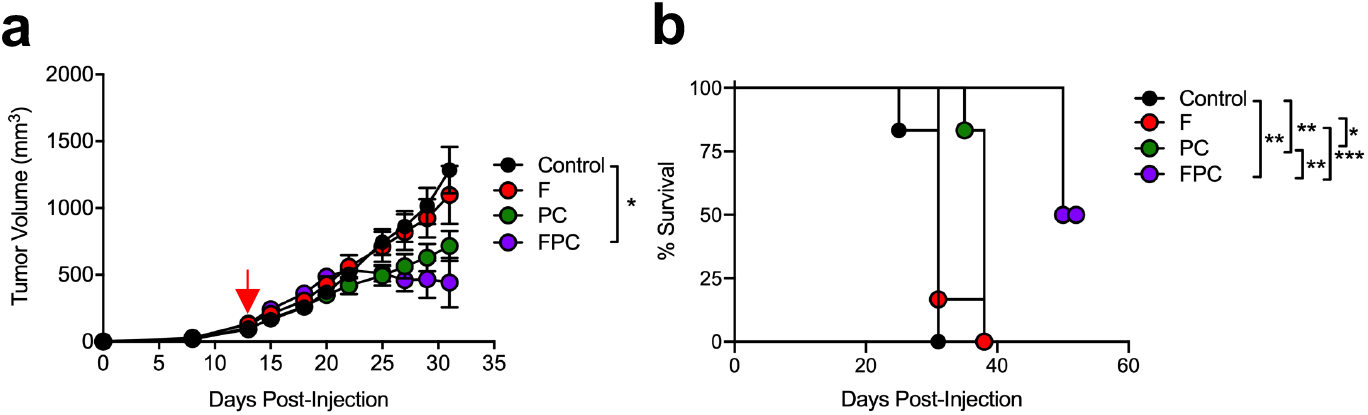
Related to Figure 1. Combination therapy with αCD40 is required for tumor regression and therapeutic efficacy. (A) Wild-type (WT) C57BL/6 mice were injected with KPCY T cell high tumor clone 2838c3 and treated with anti-PD-1 and anti-CTLA-4 (PC) on days 0,3, and 6, followed by P alone every 3 days, and a single dose of agonistic anti-CD40 (F) on day 3, alone or in combination (FPC). Mice were monitored for tumor growth (A) and survival (B). Data representative of 2 independent experiments with n=4-9 mice per group. Analysis by two-way ANOVA (A) or Mantel-Cox log-rank analysis (B), * indicates p<0.05, ** indicates p<0.01, *** indicates p<0.001.

**Extended Data Figure 2.**
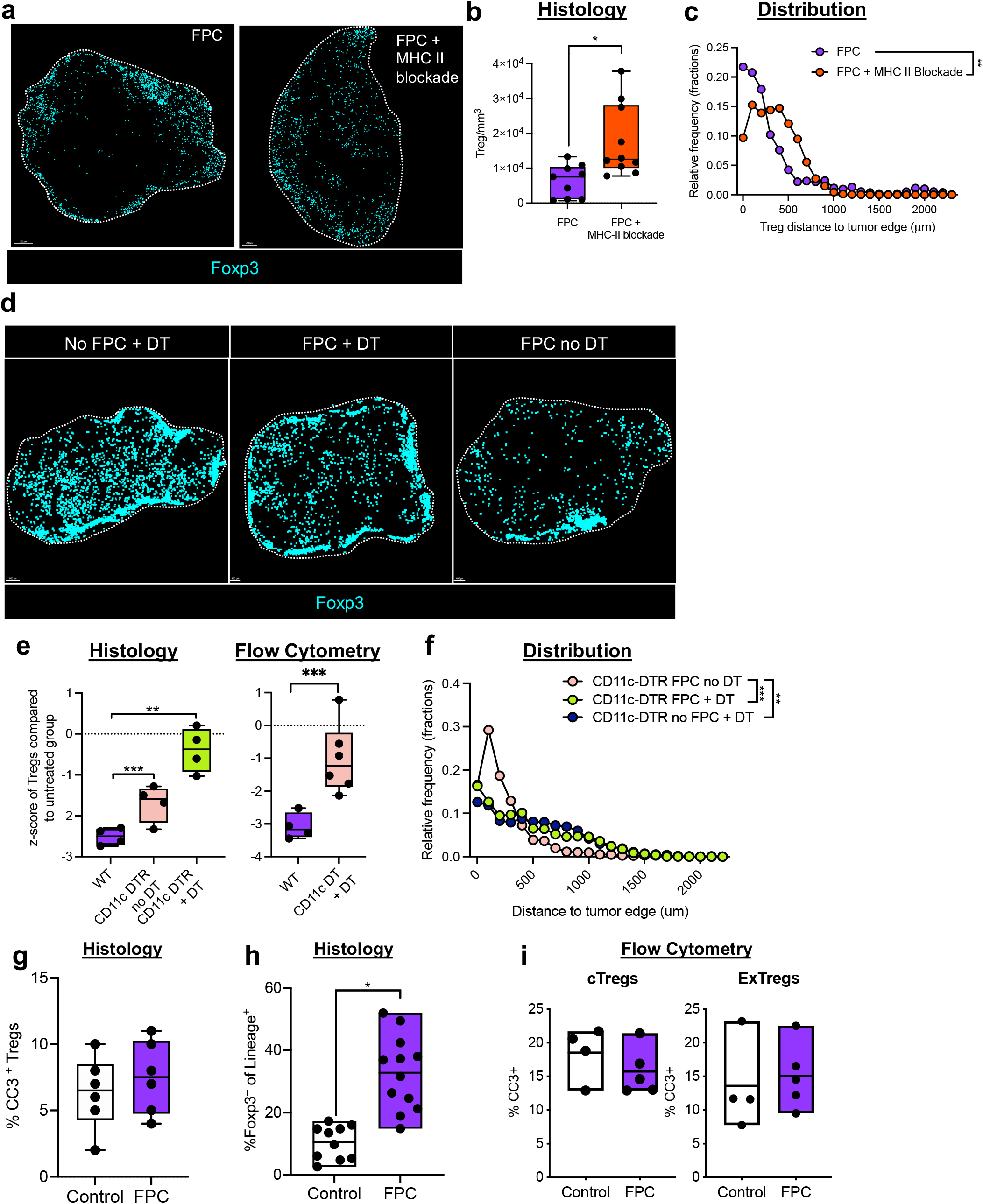
Related to Figure 2. Dendritic cells mediate αCD40-induced reduction of intratumoral Tregs. (A-C) Tumor-bearing wild-type mice were treated with FPC as described in Figure 1A, +/-the addition of MHC II blocking antibody beginning on day -2 before the start of therapy. Tumors were assessed for total numbers of Treg counts (B) and localization of Tregs (C). (D-F) CD11c DTR mice were treated as described in Fig 1A except indicated groups also received DT before assessment for Treg abundance (E) and distribution (F) in the tumor site. (G) Treg expression of cleaved caspase 3 (CC3) +/-FPC treatment in wild-type mice among live, CD45+ CD4+ T cells. (H-I) total ExTregs (tdTomato+Foxp3-) or CC3 expression among ExTregs or cTregs (tdTomato+Foxp3+) cells among live, CD45+CD4+ T cells in Foxp3 Lineage mice. Data representative of 2-5 independent experiments with n=4-9 mice per group, each symbol represents an individual mouse, horizontal lines indicate the median, the brackets of the box indicate interquartile range, and whiskers indicate the range. Scale bars for A: FPC, 300 um; FPC + MHC-II blockade, 200 um; for D: 500 um for all. Analysis by unpaired T-test (B, E, right, G-I), one-way ANOVA with Tukey’s post-test mean difference calculations to account for effect size (C, F), or one-way ANOVA (E left). For p values, * indicates p<0.05, ** indicates p<0.01, *** indicates p<0.001. For distribution plots (C, F): ** indicates mean difference of >100 um and ***>150 um.

**Extended Data Figure 3.**
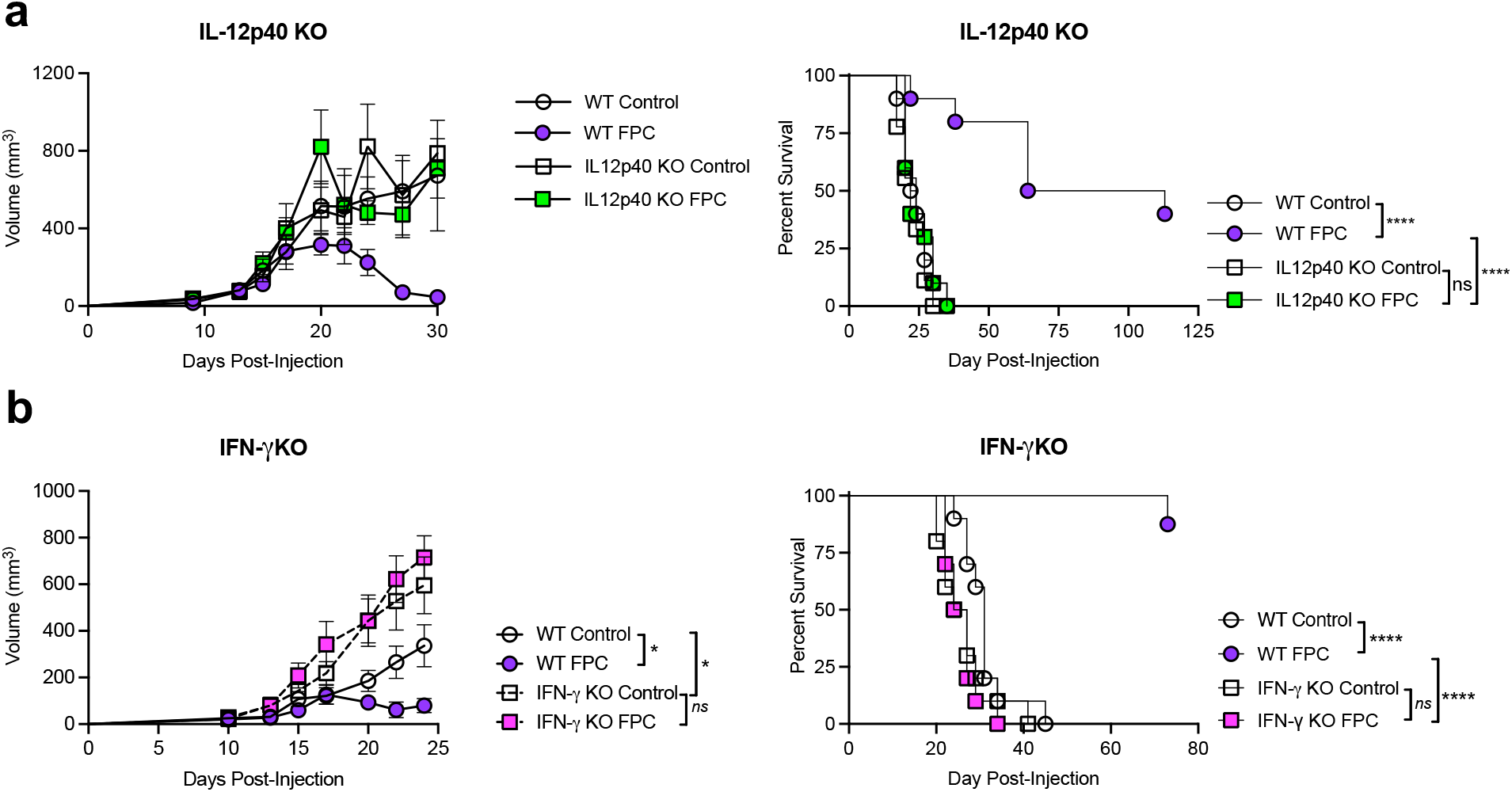
Related to Figure 3. Treg reduction is not due to cell death, and IL-12p40/IFN-*γ* axis is required for therapeutic efficacy. (A-B) Tumor-growth kinetics (right) or survival curves (left) from IL-12p40 (A) or IFN-γ (B) global knockout mice. Data representative of 2-3 independent experiments with n=4-10 mice per group, each symbol represents the mean of the group as indicated, error bars indicate SEM (A-B). Analysis by two-way ANOVA with mixed effect modeling with Tukey’s post-hoc analysis (left panels) or Mantel-Cox log-rank analysis (right panels) (D-E). * indicates p<0.05, **** indicates p<0.0001, *ns* indicates not significant.

**Extended Data Figure 4.**
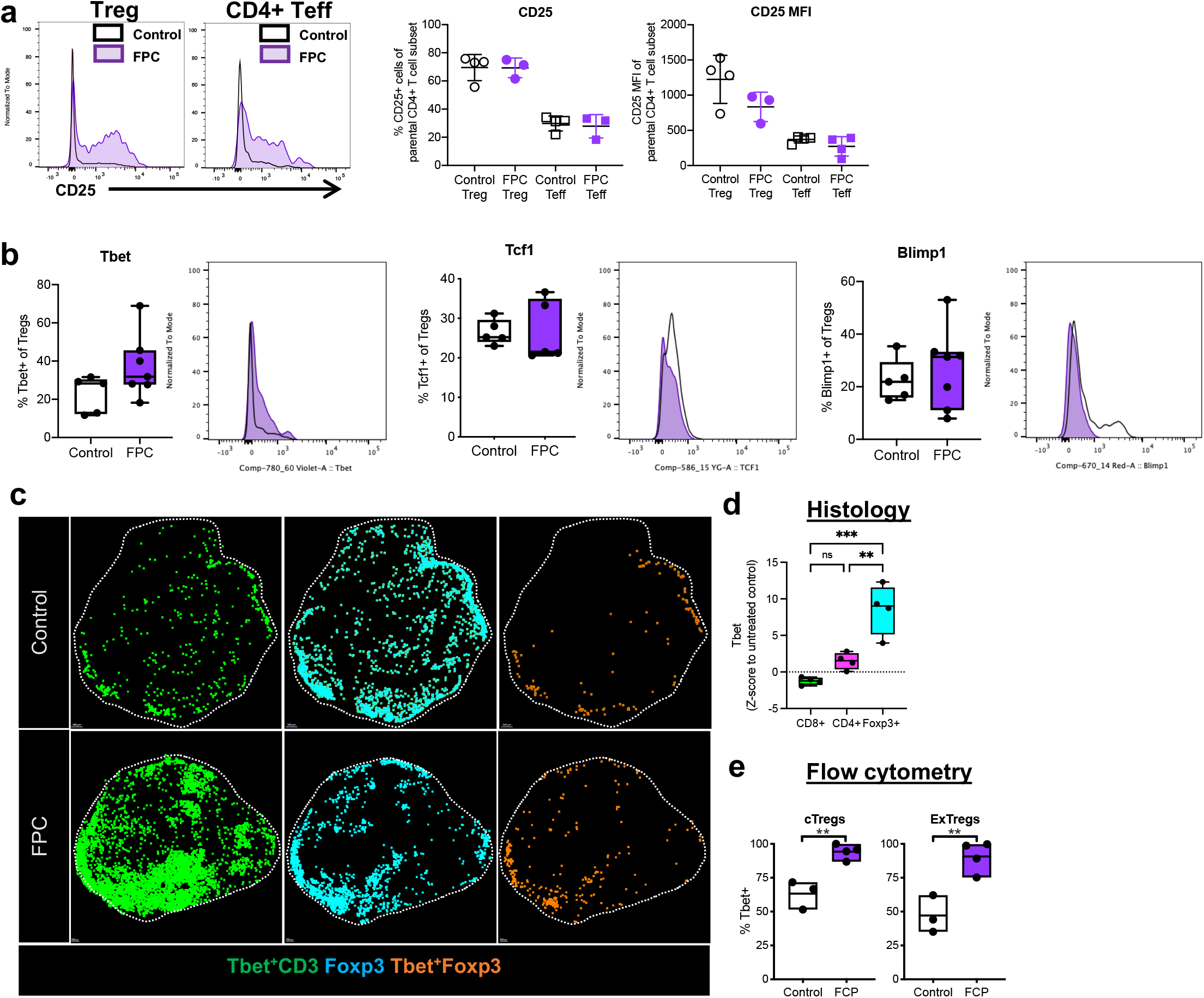
Related to Figure 4. αCD40 alters Treg transcriptional profiling. (A-B) WT mice were treated as in Fig. 1B and tumors were analyzed by intracellular flow cytometry for CD25 (A) or indicated transcription factor expression (B) among live, CD45+ CD4+ +/-Foxp3+ T cells as indicated, quantified on the left, with a representative histogram shown on the right for each marker from mice treated with vehicle Control (open) or FPC (purple). (C-D) Foxp3 and Tbet reporter mice were treated as in Fig. 1A and tumors were analyzed by tissue immunostaining (C) for expression of Tbet, Foxp3, or both (TbetFoxp3) among total CD3+ T cells, CD3+CD8+ cells, or CD4+CD3+ cells, quantified in (D). (E) Tumors were analyzed by flow cytometry for the indicated Treg subset expression of Tbet among live, CD45+CD4+ T cells. Data representative of 2-5 independent experiments with n=3-12 mice per group; each symbol represents an individual mouse, horizontal lines indicate the median, bars show interquartile range, and whiskers indicate the range (A, B, D) or bars indicate range and horizontal line indicates the mean (E). Scale bars for C: Control, 300 μm; FPC, 200 μm. Dotted line indicates tumor edge. Analysis by unpaired T-test (A, B, E), one-way ANOVA (D) with Tukey’s post-test. For p values, * indicates p<0.05, ** p<0.01, *** p<0.001, *ns* indicates not significant.

**Extended Data Figure 5.**
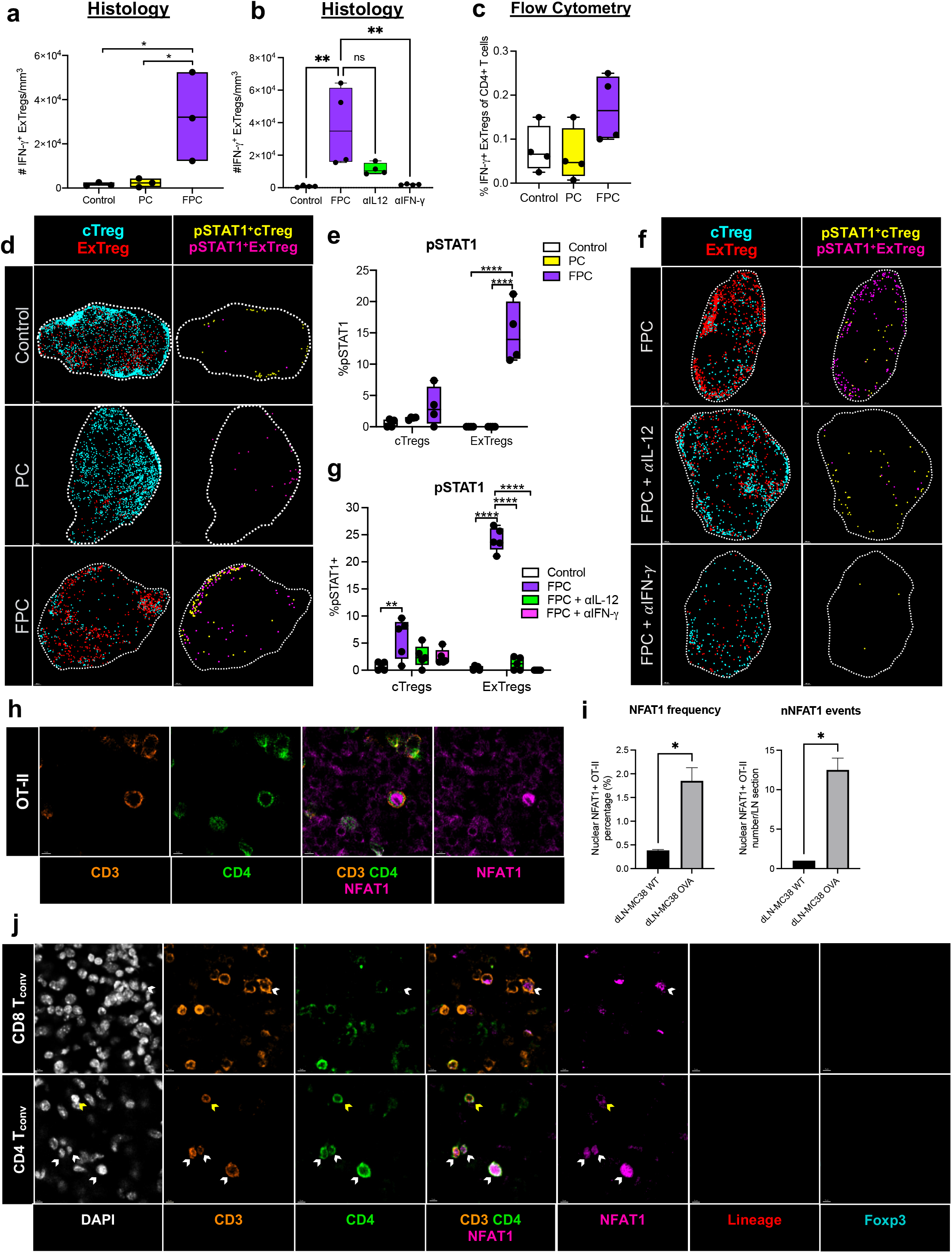
Related to Figure 4. αCD40-induced ExTregs are skewed to Th1 phenotype. (A-C) Foxp3 Lineage mice were treated as described in Fig. 3. Quantification of IFN-γ+ ExTregs by histology after treatment with vehicle control, PC, or FPC (A) or control vs. FPC +/-αIL-12p40 or αIFN-γ (B). (C) Quantification of IFN-*γ* protein by flow cytometry in control, PC, or FPC treated mice (representative of 4 experiments). (D-G) Immunofluorescent staining of pSTAT1 expression among indicated Treg subsets in Foxp3 Lineage mice after treatment with vehicle control, PC, or FPC (D) or control vs. FPC +/-αIL-12p40 or αIFN-γ (F). Quantification of pSTAT1 expression in (E) and (G). (H) MC38-OVA or MC38-WT bearing *Rag1* mice were adoptively transferred with 1×10^6^ in vitro activated OT-II cells. The tumor draining lymph node was analyzed for nuclear NFAT1 signal in OT-II cells by percentage (I, left) and number (I, right). (J) Representative tumor immunostaining for nuclear NFAT1 (white arrows) and membrane NFAT1 (yellow arrows) expression within CD8 (J, upper), and effector CD4 (non-Treg) (J, lower) conventional T cells (T_conv_) in tumor-bearing FoxP3+ Lineage mice 48 hours after FPC treatment. Data representative of 2-5 independent experiments with n=3-12 mice per group; each symbol represents an individual mouse, horizontal lines indicate the median, bar indicate range and horizontal line indicates the mean (A, B, C, E, G, I). Scale bars for D: Control, 300 μm; PC, 150 μm; FPC, 200 μm; for F: FPC and anti-IL12p40, 200 μm; anti-IFN-γ, 100 μm; for H: 5 μm; for J: 5μm. Dotted line indicates tumor edge. Analysis by one-way ANOVA (A, B, C, E, G) or unpaired T-test (I). For p values, * indicates p<0.05, ** p<0.01, *** p<0.001, *ns* indicates not significant.

## REFERENCES

1. Sharma, P., Hu-Lieskovan, S., Wargo, J. A. & Ribas, A. Primary, Adaptive, and Acquired Resistance to Cancer Immunotherapy. Cell 168, 707–723 (2017).

2. Kreiter, S. et al. Mutant MHC class II epitopes drive therapeutic immune responses to cancer. Nature 520, 692–696 (2015).

3. Alspach, E. et al. MHC-II neoantigens shape tumour immunity and response to immunotherapy. Nature 574, 696–701 (2019).

4. Ferris, S. T. et al. cDC1 prime and are licensed by CD4+ T cells to induce anti-tumour immunity. Nature 1–6 (2020) doi:10.1038/s41586-020-2611-3.

5. Morrison, A. H., Diamond, M. S., Hay, C. A., Byrne, K. T. & Vonderheide, R. H. Sufficiency of CD40 activation and immune checkpoint blockade for T cell priming and tumor immunity. Proc National Acad Sci 201918971 (2020) doi:10.1073/pnas.1918971117.

6. Śledzińska, A. et al. Regulatory T Cells Restrain Interleukin-2- and Blimp-1-Dependent Acquisition of Cytotoxic Function by CD4+ T Cells. Immunity 52, 151-166.e6 (2020).

7. Bogen, B., Fauskanger, M., Haabeth, O. A. & Tveita, A. CD4+ T cells indirectly kill tumor cells via induction of cytotoxic macrophages in mouse models. Cancer Immunol Immunother 68, 1865–1873 (2019).

8. Reed, C. M., Cresce, N. D., Mauldin, I. S., Slingluff, C. L. & Olson, W. C. Vaccination with Melanoma Helper Peptides Induces Antibody Responses Associated with Improved Overall Survival. Clin Cancer Res 21, 3879–3887 (2015).

9. Rouhani, S. J. et al. Roles of lymphatic endothelial cells expressing peripheral tissue antigens in CD4 T-cell tolerance induction. Nat Commun 6, 6771 (2015).

10. Hingorani, S. R. et al. Trp53R172H and KrasG12D cooperate to promote chromosomal instability and widely metastatic pancreatic ductal adenocarcinoma in mice. Cancer Cell 7, 469–483 (2005).

11. Li, J. et al. Tumor Cell-Intrinsic Factors Underlie Heterogeneity of Immune Cell Infiltration and Response to Immunotherapy. Immunity 49, 178-193.e7 (2018).

12. Morrison, A. H., Diamond, M. S., Hay, C. A., Byrne, K. T. & Vonderheide, R. H. Sufficiency of CD40 activation and immune checkpoint blockade for T cell priming and tumor immunity. Proc National Acad Sci 201918971 (2020) doi:10.1073/pnas.1918971117.

13. Diehl, L. et al. CD40 activation in vivo overcomes peptide-induced peripheral cytotoxic T-lymphocyte tolerance and augments anti-tumor vaccine efficacy. Nat Med 5, 774–779 (1999).

14. Sotomayor, E. M. et al. Conversion of tumor-specific CD4+ T-cell tolerance to T-cell priming through in vivo ligation of CD40. Nat Med 5, 780–787 (1999).

15. French, R. R., Chan, H. T. C., Tutt, A. L. & Glennie, M. J. CD40 antibody evokes a cytotoxic T-cell response that eradicates lymphoma and bypasses T-cell help. Nat Med 5, 548–553 (1999).

16. Ferris, S. T. et al. cDC1 prime and are licensed by CD4+ T cells to induce anti-tumour immunity. Nature 1–6 (2020) doi:10.1038/s41586-020-2611-3.

17. Byrne, K. T. & Vonderheide, R. H. CD40 Stimulation Obviates Innate Sensors and Drives T Cell Immunity in Cancer. Cell Reports 15, 2719–2732 (2016).

18. Winograd, R. et al. Induction of T-cell Immunity Overcomes Complete Resistance to PD-1 and CTLA-4 Blockade and Improves Survival in Pancreatic Carcinoma. Cancer Immunol Res 3, 399–411 (2015).

19. Gerner, M. Y., Kastenmuller, W., Ifrim, I., Kabat, J. & Germain, R. N. Histo-Cytometry: A Method for Highly Multiplex Quantitative Tissue Imaging Analysis Applied to Dendritic Cell Subset Microanatomy in Lymph Nodes. Immunity 37, 364–376 (2012).

20. Bulliard, Y. et al. Activating Fc γ receptors contribute to the antitumor activities of immunoregulatory receptor-targeting antibodies. J Exp Medicine 210, 1685–1693 (2013).

21. Simpson, T. R. et al. Fc-dependent depletion of tumor-infiltrating regulatory T cells co-defines the efficacy of anti–CTLA-4 therapy against melanoma. J Exp Med 210, 1695–1710 (2013).

22. Tang, T. et al. Molecular basis and therapeutic implications of CD40/CD40L immune checkpoint. Pharmacol Therapeut 219, 107709 (2020).

23. Lin, J. H. et al. Type 1 conventional dendritic cells are systemically dysregulated early in pancreatic carcinogenesis. J Exp Medicine 217, e20190673 (2020).

24. Hildner, K. et al. Batf3 Deficiency Reveals a Critical Role for CD8α+ Dendritic Cells in Cytotoxic T Cell Immunity. Science 322, 1097–1100 (2008).

25. Hor, J. L. et al. Spatiotemporally Distinct Interactions with Dendritic Cell Subsets Facilitates CD4+ and CD8+ T Cell Activation to Localized Viral Infection. Immunity 43, 554–565 (2015).

26. Kastenmüller, W., Gerner, M. Y. & Germain, R. N. The in situ dynamics of dendritic cell interactions. Eur J Immunol 40, 2103–2106 (2010).

27. Rubtsov, Y. P. et al. Stability of the Regulatory T Cell Lineage in Vivo. Science 329, 1667–1671 (2010).

28. Hsieh, C.-S. et al. Development of TH1 CD4+ T Cells Through IL-12 Produced by Listeria-Induced Macrophages. Science 260, 547–549 (1993).

29. Oldenhove, G. et al. Decrease of Foxp3+ Treg Cell Number and Acquisition of Effector Cell Phenotype during Lethal Infection. Immunity 31, 772–786 (2009).

30. Alspach, E., Lussier, D. M. & Schreiber, R. D. Interferon γ and Its Important Roles in Promoting and Inhibiting Spontaneous and Therapeutic Cancer Immunity. Csh Perspect Biol 11, a028480 (2019).

31. Gocher, A. M., Workman, C. J. & Vignali, D. A. A. Interferon-γ: teammate or opponent in the tumour microenvironment? Nat Rev Immunol 22, 158–172 (2022).

32. Thornton, A. M. et al. Helios+ and Helios− Treg subpopulations are phenotypically and functionally distinct and express dissimilar TCR repertoires. Eur J Immunol 49, 398–412 (2019).

33. Thornton, A. M. & Shevach, E. M. Helios: still behind the clouds. Immunology 158, 161–170 (2019).

34. Thornton, A. M. et al. Expression of Helios, an Ikaros Transcription Factor Family Member, Differentiates Thymic-Derived from Peripherally Induced Foxp3+ T Regulatory Cells. J Immunol 184, 3433–3441 (2010).

35. Oldenhove, G. et al. Decrease of Foxp3+ Treg Cell Number and Acquisition of Effector Cell Phenotype during Lethal Infection. Immunity 31, 772–786 (2009).

36. Pilato, M. D. et al. Targeting the CBM complex causes Treg cells to prime tumours for immune checkpoint therapy. Nature 570, 112–116 (2019).

37. Daniel, V., Sadeghi, M., Wang, H. & Opelz, G. CD4+CD25+Foxp3+IFN-γ+ human induced T regulatory cells are induced by interferon-γ and suppress alloresponses nonspecifically. Hum Immunol 72, 699–707 (2011).

38. Koch, M. A. et al. T-bet controls regulatory T cell homeostasis and function during type-1 inflammation. Nat Immunol 10, 595–602 (2009).

39. Teijaro, J. R. et al. Persistent LCMV Infection Is Controlled by Blockade of Type I Interferon Signaling. Science 340, 207–211 (2013).

40. Wherry, E. J. & Kurachi, M. Molecular and cellular insights into T cell exhaustion. Nat Rev Immunol 15, 486–499 (2015).

41. Crawford, A. et al. Molecular and Transcriptional Basis of CD4+ T Cell Dysfunction during Chronic Infection. Immunity 40, 289–302 (2014).

42. Wilson, E. B. et al. Blockade of Chronic Type I Interferon Signaling to Control Persistent LCMV Infection. Science 340, 202–207 (2013).

43. Hu, X. & Ivashkiv, L. B. Cross-regulation of Signaling Pathways by Interferon-γ: Implications for Immune Responses and Autoimmune Diseases. Immunity 31, 539–550 (2009).

44. Stark, G. R. How cells respond to interferons revisited: From early history to current complexity. Cytokine Growth F R 18, 419–423 (2007).

45. Macian, F. & Maclian. NFAT proteins: key regulators of T-cell development and function. Nat Rev Immunol 5, 472–484 (2005).

46. Marangoni, F. et al. The Transcription Factor NFAT Exhibits Signal Memory during Serial T Cell Interactions with Antigen-Presenting Cells. Immunity 38, 237–249 (2013).

47. Marangoni, F. et al. Expansion of tumor-associated Treg cells upon disruption of a CTLA-4-dependent feedback loop. Cell 184, 3998-4015.e19 (2021).

48. Pesic, M. et al. 2-photon imaging of phagocyte-mediated T cell activation in the CNS. J Clin Invest 123, 1192–1201 (2013).

49. Smigiel, K. S. et al. CCR7 provides localized access to IL-2 and defines homeostatically distinct regulatory T cell subsets. J Exp Med 211, 121–136 (2014).

50. Sawant, D. V. & Vignali, D. A. A. Once a Treg, always a Treg? Immunol Rev 259, 173–191 (2014).

51. Fife, B. T. et al. Interactions between PD-1 and PD-L1 promote tolerance by blocking the TCR–induced stop signal. Nat Immunol 10, 1185–1192 (2009).

52. Tadokoro, C. E. et al. Regulatory T cells inhibit stable contacts between CD4+ T cells and dendritic cells in vivo. J Exp Medicine 203, 505–511 (2006).

53. Marangoni, F. et al. Expansion of tumor-associated Treg cells upon disruption of a CTLA-4-dependent feedback loop. Cell 184, 3998-4015.e19 (2021).

54. Kawabe, T. et al. The immune responses in CD40-deficient mice: Impaired immunoglobulin class switching and germinal center formation. Immunity 1, 167–178 (1994).

55. Dalton, D. K. et al. Multiple Defects of Immune Cell Function in Mice with Disrupted Interferon-γ Genes. Science 259, 1739–1742 (1993).

56. Huang, S. et al. Immune Response in Mice that Lack the Interferon-γ Receptor. Science 259, 1742–1745 (1993).

57. Magram, J. et al. IL-12-Deficient Mice Are Defective in IFNγ Production and Type 1 Cytokine Responses. Immunity 4, 471–481 (1996).

58. Jung, S. et al. In Vivo Depletion of CD11c+ Dendritic Cells Abrogates Priming of CD8+ T Cells by Exogenous Cell-Associated Antigens. Immunity 17, 211–220 (2002).

59. Kim, J. M., Rasmussen, J. P. & Rudensky, A. Y. Regulatory T cells prevent catastrophic autoimmunity throughout the lifespan of mice. Nat Immunol 8, 191–197 (2007).

60. Rubtsov, Y. P. et al. Stability of the Regulatory T Cell Lineage in Vivo. Science 329, 1667–1671 (2010).

61. Madisen, L. et al. A robust and high-throughput Cre reporting and characterization system for the whole mouse brain. Nat Neurosci 13, 133–140 (2010).

62. Reinhardt, R. L., Liang, H.-E. & Locksley, R. M. Cytokine-secreting follicular T cells shape the antibody repertoire. Nat Immunol 10, 385–393 (2009).

63. Radtke, A. J. et al. IBEX: A versatile multiplex optical imaging approach for deep phenotyping and spatial analysis of cells in complex tissues. Proc National Acad Sci 117, 33455–33465 (2020).

